# cpn60 barcode sequences accurately identify newly defined genera within the *Lactobacillaceae*

**DOI:** 10.1101/2021.02.24.432354

**Authors:** Ishika Shukla, Janet E. Hill

**Affiliations:** Department of Veterinary Microbiology, University of Saskatchewan, Saskatoon, Saskatchewan, Canada

**Keywords:** *Lactobacillaceae*, cpn60, hsp60, groEL, barcoding, taxonomy, *Lactobacillus*, metabarcoding

## Abstract

The cpn60 barcode sequence is established as an informative target for microbial species identification. Applications of cpn60 barcode sequencing are supported by the availability of “universal” PCR primers for its amplification and a curated reference database of cpn60 sequences, cpnDB. A recent reclassification of lactobacilli involving the definition of 23 new genera provided an opportunity to update cpnDB and to determine if the cpn60 barcode could be used for accurate identification of species consistent with the new framework. Analysis of 275 cpn60 sequences representing 258/269 of the validly named species in *Lactobacillus, Paralactobacillus* and the 23 newer genera showed that cpn60-based sequence relationships were consistent with the whole-genome-based phylogeny. Aligning or mapping full length barcode sequences or a 150 bp subsequence resulted in accurate and unambiguous species identification in almost all cases. Taken together, our results show that the combination of available reference sequence data, “universal” barcode amplification primers, and the inherent sequence diversity within the cpn60 barcode make it a useful target for the detection and identification of lactobacilli as defined by the latest taxonomic framework.

**Significance and Impact of the Study:** The genus *Lactobacillus* recently underwent a major reorganization resulting in the definition of 23 new genera. Lactobacilli are widespread in environmental and host-associated microbiomes and are exploited in food and biotechnology applications, making methods for their accurate identification desirable. Here we show that the combination of a reference sequence database, “universal” barcode amplification primers, and the inherent sequence diversity within the cpn60 barcode make it a useful target for the detection and identification of lactobacilli as defined by the latest taxonomic framework.

## INTRODUCTION

Amplicon sequencing of informative taxonomic marker genes is a mainstay of identification of microbial species either in isolation or in complex microbial communities. The region of the cpn60 gene (also known as groEL or hsp60) corresponding to nucleotides 271-822 of the *Escherichia coli* cpn60 gene meets the criteria for a barcode sequence for bacteria (Links *et al.*, 2012). The species-level resolution provided by the cpn60 barcode has led to its use in characterization of a wide range of microbial communities (Dumonceaux *et al.*, 2006b; Oliver, Hamelin & Hintz, 2008; Pratt *et al.*, 2012; Freitas *et al.*, 2018; King *et al.*, 2019; Xie *et al.*, 2019), and in taxonomic and phylogenetic studies (Carlier, Bonne & Bedora-Faure, 2006; Hill *et al.*, 2006a; Minana-Galbis *et al.*, 2009; Sakamoto & Ohkuma, 2010). It was established recently that 150 bp from the 5’ end of the barcode is generally sufficient for species level identification (Vancuren *et al.*, 2020). “Universal” PCR primers are available for its amplification (Hill, Town & Hemmingsen, 2006b) and a reference database of cpn60 sequences, cpnDB, has supported its application to microbial species identification since 2004 (www.cpndb.ca) (Vancuren & Hill, 2019). The goal of cpnDB is to provide high quality, manually curated records representing the breadth of prokaryotic taxonomy, with an emphasis on type strains. Revisions to taxonomy require corresponding database updates. These amendments also offer opportunities to evaluate the ability of cpn60 barcode sequences to resolve species, and to determine if cpn60 barcode relationships are consistent with new taxonomy frameworks, such as was recently demonstrated for *Gardnerella* spp. (Hill & Albert, 2019).

*Lactobacillus* spp. are widespread in environmental and host-associated microbiomes and they have been exploited in food and biotechnology applications (Bosma, Forster & Nielsen, 2017; Sauer *et al.*, 2017). Ongoing discovery of new taxa led to the rapid expansion of the genus to include over 250 validly named species by 2019. Various regions of the cpn60 gene have been used to identify and distinguish *Lactobacillus* spp. and other closely related lactic acid bacteria either in isolation or in metagenomic samples because it has been established that the sequence is generally more variable among species than 16S rRNA gene sequences (Blaiotta *et al.*, 2008; Haakensen, Pittet & Ziola, 2011; Koirala *et al.*, 2015; Freitas *et al.*, 2018; Xie *et al.*, 2019; Yang *et al.*, 2019; Damé-Teixeira *et al.*, 2020).

Recently, an examination of the *Lactobacillus* genus and related taxa in the family *Lactobacillaceae* led to a proposal to reclassify many *Lactobacillus* spp. into 23 new genera based primarily on analysis of whole genome sequences (Zheng *et al.*, 2020). Given their importance and prevalence, robust molecular tools for the identification and discrimination of *Lactobacillaceae* are desirable. Inspired by the new taxonomic framework for *Lactobacillus,* the objectives of this study were to update *Lactobacillaceae* records in the cpnDB database, determine if cpn60 sequence relationships are consistent with the newly designated genera, and to assess the accuracy of identification of species based on alignment and mapping of barcode sequences using methods commonly employed in metagenomic studies.

## RESULTS & DISCUSSION

### Curation of cpnDB

Of the 286 taxa identified for inclusion in the study, cpn60 sequences were available for all but 11 (Table S1). For these 11 taxa, no whole genome sequence was available as of December 2020 but the species is validly published and documented in the LPSN database. The missing taxa included seven genera: *Fructilactobacillus* (2 species), *Furfurilactobacillus* (1 species), *Lacticaseibacillus* (1 species), *Lactobacillus* (3 species), *Ligilactobacillus* (1 species), *Limosilactobacillus* (2 species), and *Liquorilactobacillus* (1 species). Other representatives of each of these genera, however, were available (Table 1). Following the update to cpnDB to account for new genus names and new taxa, the database now contains cpn60 sequences for at least one representative of 258/269 of the species in *Lactobacillus, Paralactobacillus* and the 23 newer genera that were validly named at the time of the study. If and when sequences become available for the currently unrepresented taxa they will be incorporated as part of the ongoing curation of cpnDB. We encourage interested users to notify us in cases where publicly available sequences are not available through cpnDB.

**Table 1.**
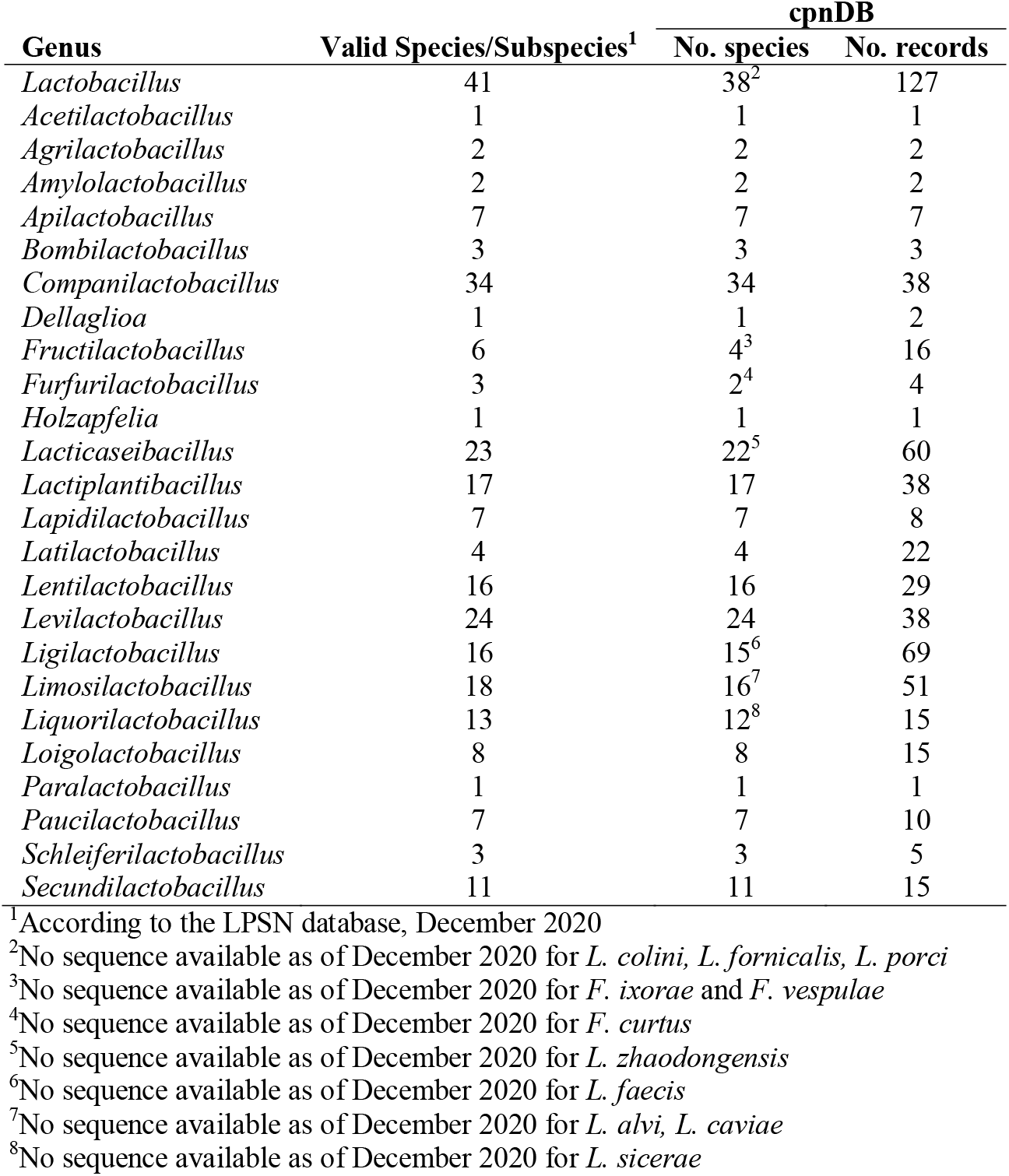
Numbers of valid species/subspecies, cpnDB records and species represented for *Lactobacillus, Paralactobacillus* and 23 newly defined genera.

### cpn60 sequence alignments and comparisons

All of the cpn60 barcode sequences included in this study were 552 bp in length, making multiple sequence alignment trivial. The cpn60 barcodes had a median G+C content of 0.41 and these values were not significantly different than those reported for the corresponding whole genome sequences (Table S1, Mann-Whitney, p = 0.842).

The definition of new genera with the family *Lactobacillaceae* was based largely on whole genome sequence comparisons (Zheng *et al.*, 2020). Previous studies have shown that cpn60 barcode sequences can be used as predictors of whole genome sequence relationships across a range of taxa (Verbeke *et al.*, 2011; Schellenberg *et al.*, 2016), which has resulted in the use of this target (also known as hsp60 or groEL) for identification and discrimination of related bacteria in various settings, addressing phylogenetic questions and the designation of new species including lactobacilli (Hiu *et al.*, 1984; Brousseau *et al.*, 2001; Bringel *et al.*, 2005; Pantucek *et al.*, 2005; Carlier *et al.*, 2006; Hill *et al.*, 2006a; Claesson, van Sinderen & O’Toole, 2008; Minana-Galbis *et al.*, 2009; Sakamoto, Suzuki & Benno, 2010; Sakamoto *et al.*, 2018; Sakamoto & Ohkuma, 2010; Haakensen *et al.*, 2011; Hill & Albert, 2019; Muirhead *et al.*, 2019). To determine if cpn60 barcode sequence relationships corresponded to the new taxonomic framework for lactobacilli, a maximum likelihood tree was constructed from 275 cpn60 barcode sequences (Figure 1). Genera formed discrete clusters with a few exceptions. *Secundilactobacillus malefermentans* and *S. orzae* were distant from the other members of the genus; a similar observation was made for *Ligilactobacillus* where five species (*acidipiscis, salitolerans, pobuzihii, agilis* and *equi*) were well separated from the rest of the genus. This was not unexpected since these genera were also described as “non-exclusive” by Zheng et al. since the lower *intra*-genus cAAI (pairwise amino acid identity of conserved genes) was lower than the highest *inter*-genus cAAI (Zheng *et al.*, 2020). *Lactobacillus bombintestini* was distant from the other *Lactobacillus* spp. and clustered with *Apilactobacillus.* The publication of *L. bombintestini* occurred very recently, and concurrently with the proposed reclassification of lactobacilli (Heo *et al.*, 2020). Its cpn60 sequence clearly clusters within the new genus *Apilactobacillus* with other bee-associated species (Figure 1) and suggests that this species may soon be reclassified.

**Figure 1.**
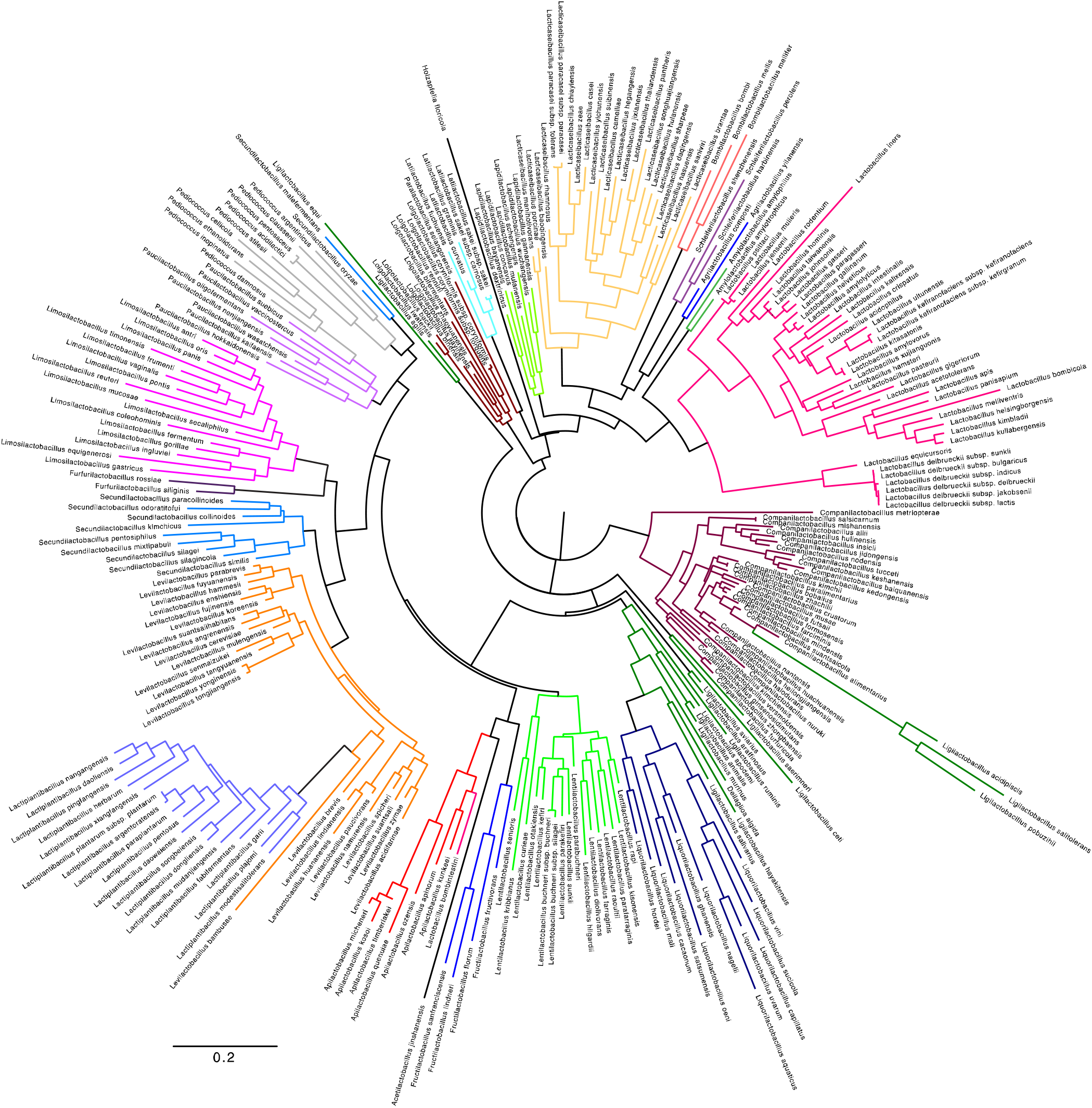
Maximum likelihood tree of 275 *Lactobacillaceae* cpn60 barcode sequences was inferred by PHYML using the GTR+G+I model. The tree is displayed with midpoint root. Members of the same genus are indicated by branch colour.

Overall pairwise percent nucleotide identity values among all *Lactobacillaceae* sequences ranged from 64-100% (median 76.8%) (Figure 2). The only pairs of identical sequences were those from two pairs of subspecies of *Lactobacillus delbrueckii* (subspecies *jakobsenii* and *lactis*, and subspecies *bulgaricus* and *sunkii*), and two pairs of species of *Companilactobacillus* (*salsicarnum* and *michanensis*, *bobalius* and *paralimentarius*). Comparison of the intra-genus pairwise similarities revealed median intra-genus identities ranging from 79.2% for *Ligilactobacillus* to 90.9% for *Latilactobacillus* (Figure 3). This pattern of *intra*-genus pairwise sequence identities for cpn60 mirrors the results of comparison of conserved proteins from the complete genomes (Zheng *et al.*, 2020).

**Figure 2.**
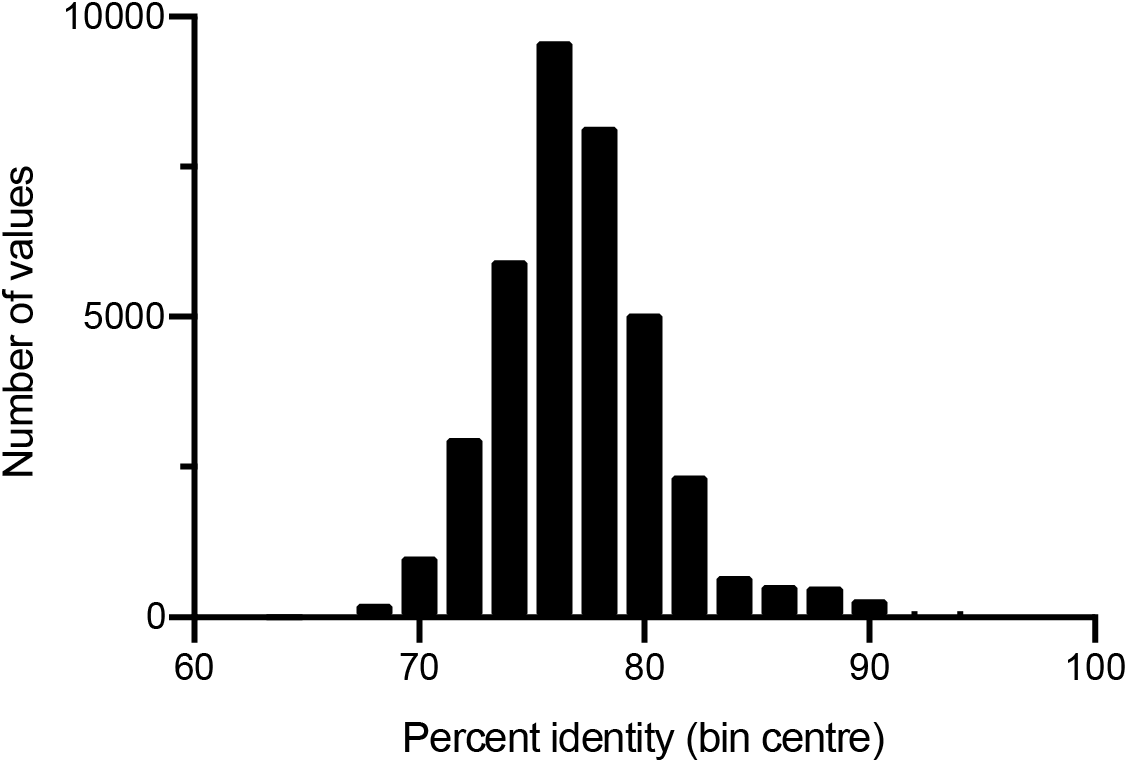
Distribution of pairwise nucleotide sequence identity for 275 cpn60 sequences (37,675 comparisons).

**Figure 3.**
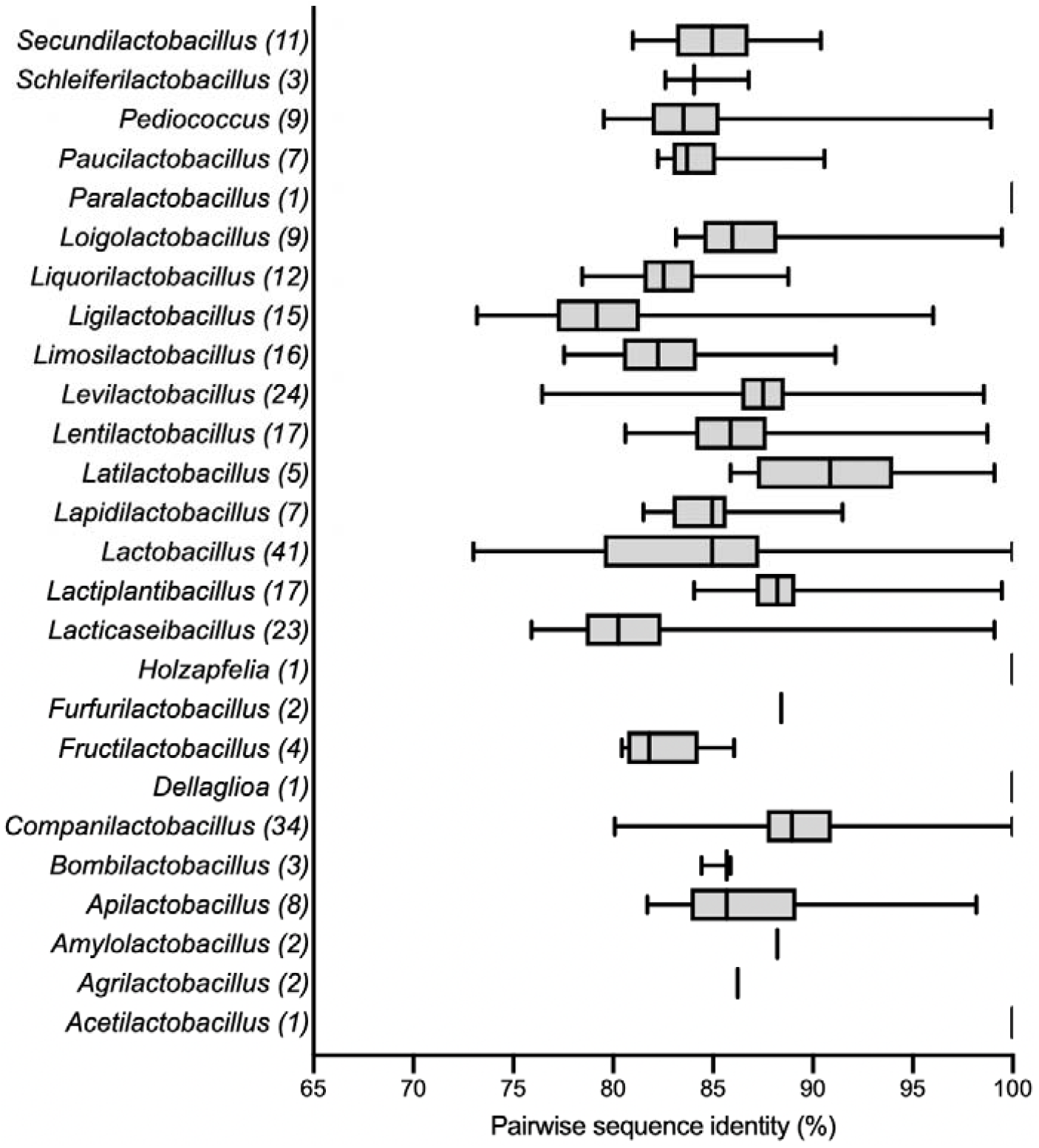
Intra-genus pairwise nucleotide sequence identities. Number of sequences in each genus is indicated in parentheses. Boxes indicated the 25^th^ to 75^th^ percentile values and median; whiskers show minimum to maximum. The trivial value of 100% is shown for genera with only one representative.

### Accuracy of cpn60-based sequence identification

To determine if the sequence diversity within the cpn60 barcode sequence was sufficient to provide unambiguous identification, three approaches were used based on the methods commonly used to compare cpn60 barcode sequences from isolates or metagenomic samples to reference sequences. Each experiment was performed with both the entire 552 bp of the barcode region or a 150 bp subsequence (nucleotides 1-150 of the barcode). In each case, the reference database was comprised of the 275 study sequences.

Query sequences were aligned to the reference data set using FASTA36 and BLASTn (Figure 4). Since alignments generally covered the entire length of the query, the percent identity for each of the top five results returned were evaluated. For both query lengths, using either alignment method, there was a significant drop in sequence identity between the first hit (100% identity to self) and the next best match (median percent identity ~90% for both query lengths by either alignment method) (Wilcoxon test, p<0.0001). For 75% of queries, the first non-self hit was <95% identical, making identification and discrimination of these members of the *Lactobacillaceae* relatively simple, as has been demonstrated previously for lactobacilli in isolation (Blaiotta *et al.*, 2008; Haakensen *et al.*, 2009) or within host-associated microbial communities containing lactobacilli such as the gastrointestinal and vaginal microbiota where species-level identification is achieved routinely with cpn60 barcode sequences (Dumonceaux *et al.*, 2006a; Chaban *et al.*, 2014; Freitas *et al.*, 2018).

**Figure 4.**
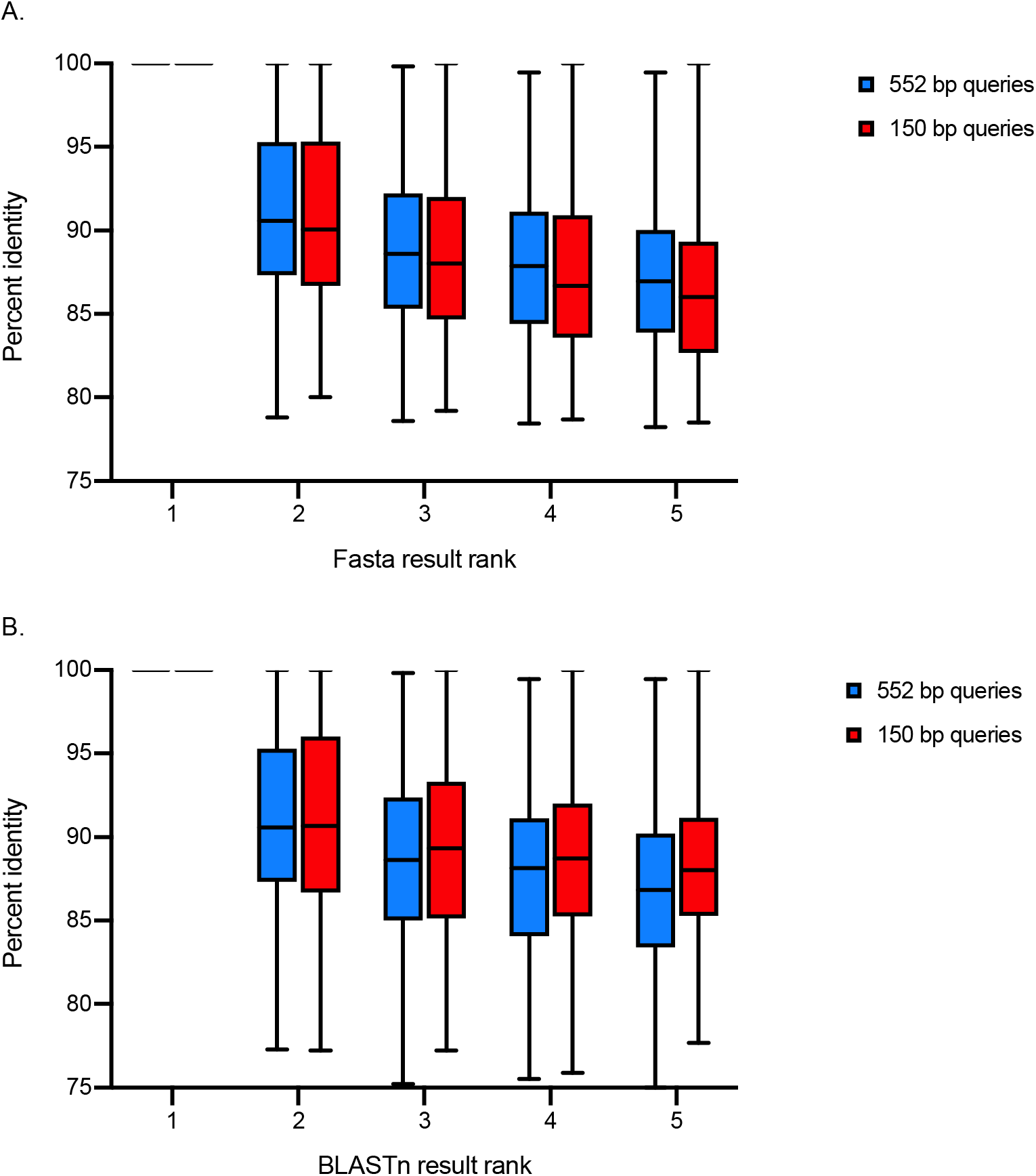
Percent identity for top five hits for each 552 bp and 150 bp query aligned to the reference data set using (A) FASTA36 or (B) BLASTn. Whiskers indicate minimum to maximum values. The top ranked result for each query is itself (all 100% identity).

For microbial community analysis based on high throughput sequencing of barcode sequences (“metabarcoding”), relatively short read-lengths and large data sets are additional challenges for species identification. Rapid read mapping approaches using tools such as Minimap2 that are based on the frequency of short subsequences (kmers) are commonly utilized in these cases since they are more efficient and rapid than alignment-based methods for comparisons of very large numbers of queries with large databases (Hatem *et al.*, 2013). Mapping of queries with Minimap2 returned only one result (itself) in the large majority of cases for both 552 bp (236/275, 85.8%) and 150 bp queries (233/275, 84.7%) (Figure 5). The demonstration that the 150 bp subsequence is generally sufficient for species-level identification is consistent with previous observations (Vancuren *et al.*, 2020) and is an advantage for rapid and economical high throughput amplicon sequencing of metagenomic samples.

**Figure 5.**
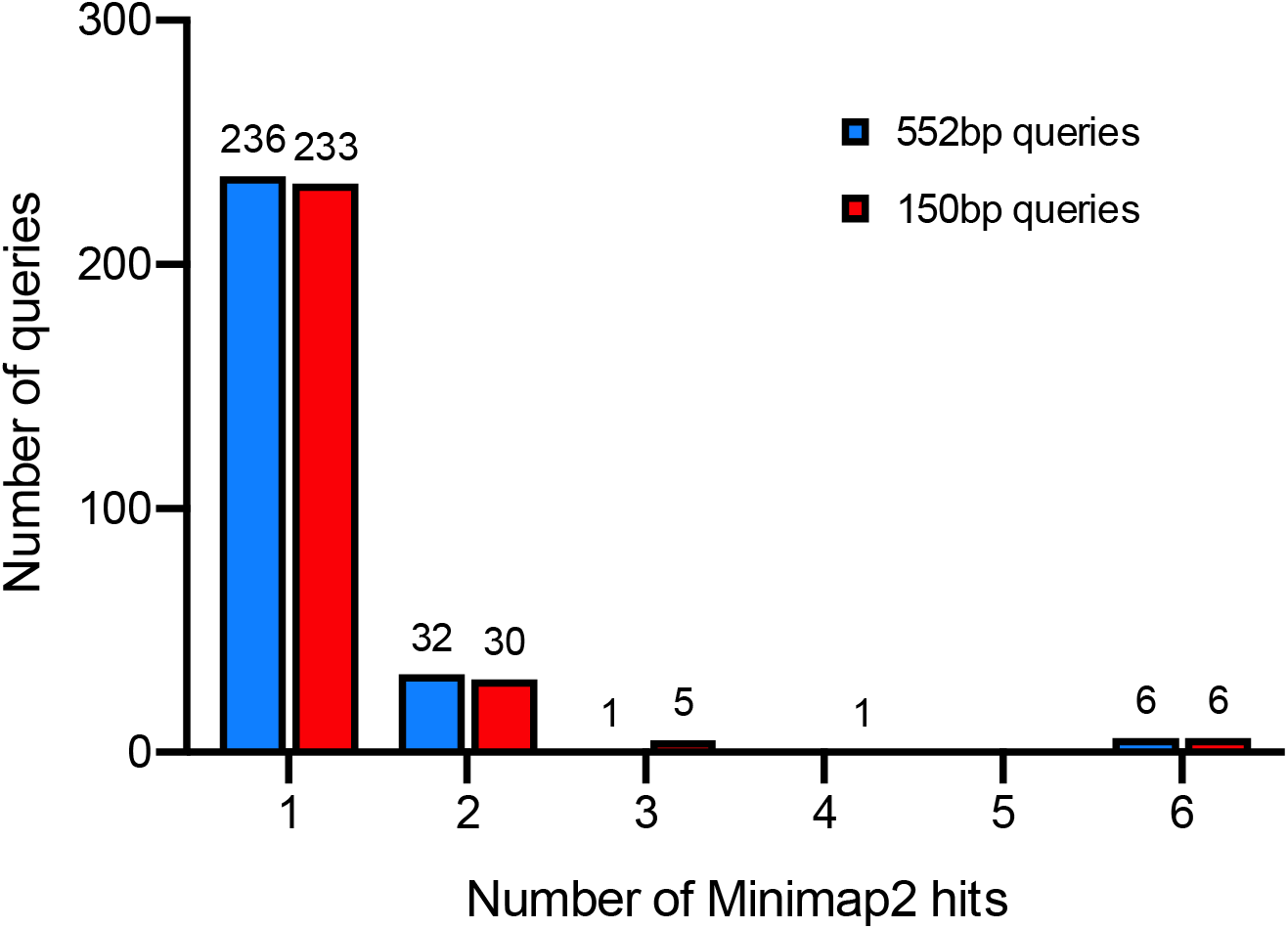
Number of results returned for 552 bp or 150 bp query sequences using Minimap2 mapping.

Taken together, our results show that the combination of available reference sequence data, “universal” barcode amplification primers, and the inherent sequence diversity within the cpn60 barcode make it a useful target for the detection and identification of lactobacilli as defined by the latest taxonomic framework.

## MATERIALS AND METHODS

### Retrieval of sequence data and update of cpnDB

A list of 281 taxa and associated genome sequences was derived from Zheng et al. (2020), and supplemented with five more recently defined *Lactobacillus* spp. (Bai *et al.*, 2020; Liu & Gu, 2020) for a final list of 286 validly published species and subspecies. In cases where the species was already represented in cpnDB (http://www.cpndb.ca), the genus name was updated to reflect the new name. If the species was not already included in cpnDB, the cpn60 sequence (full-length and barcode region) was extracted from the genome (if available) and a new record was created. Type strains and strain name synonyms for type strains were identified using the List of Prokaryotes with Standing in Nomenclature database (LPSN, https://lpsn.dsmz.de)(Parte *et al.*, 2020). A list of species, cpnDB accession numbers and Genbank accession numbers for the source genomes is provided in Table S1.

### Sequence analysis, alignments and trees

Multiple sequence alignments were conducted with CLUSTALw using default settings. Pairwise sequence identities were calculated in MegaX. Maximum likelihood trees were constructed using PhyML-SMS (Guindon *et al.*, 2010; Lefort, Longueville & Gascuel, 2017) with the GTR+G+I model. Trees were visualized and edited in FigTree version 1.4.4. G+C content was calculated using *geecee* in the EMBOSS package (Rice, Longden & Bleasby, 2000).

### Assessment of accuracy of species identification

The entire set of cpn60 barcode sequences was used to create databases for BLASTn (Altschul *et al.*, 1997), FASTA (Pearson & Lipman, 1988) and Minimap2 (Li, 2018). Each 552 bp barcode sequence and the corresponding initial 150 bp subsequence were then used as queries. For BLASTn and FASTA, the top five hits were recorded. For minimap2 the number of mapping events for each query was recorded. Differences in scores or number of mapping events were assessed using Kruskal-Wallis followed by pairwise Mann-Whitney or Wilcoxon tests if significant differences were detected. Statistical analysis was performed in GraphPad Prism 9.

## ACKNOWLEDGEMENTS

The authors are grateful to members of the Hill Lab for helpful discussions. No thanks to COVID-19. This work was supported by a Discovery Grant from the Natural Sciences and Engineering Research Council of Canada to JEH. IS was partially supported by an undergraduate research scholarship from the Western College of Veterinary Medicine.

## SUPPORTING INFORMATION

**Table S1**. 275 Lactobacillaceae species included in the study with Genbank and cpnDB accession numbers.

## REFERENCES

Altschul, S.F., Madden, T.L., Schaffer, A.A., Zhang, J., Zhang, Z., Miller, W. and Lipman, D.J. (1997) Gapped BLAST and PSI-BLAST: a new generation of protein database search programs. Nucleic Acids Research. 25, 3389–402.

Bai, L., Paek, J., Shin, Y., Park, H.-Y. and Chang, Y.H. (2020) *Lentilactobacillus kribbianus* sp. nov., isolated from the small intestine of a mini pig. International Journal of Systematic and Evolutionary Microbiology. 70, 6476–6481.

Blaiotta, G., Fusco, V., Ercolini, D., Aponte, M., Pepe, O. and Villani, F. (2008) *Lactobacillus* strain diversity based on partial hsp60 gene sequences and design of PCR-restriction fragment length polymorphism assays for species identification and differentiation. Applied and Environmental Microbiology. 74, 208–15.

Bosma, E.F., Forster, J. and Nielsen, A.T. (2017) Lactobacilli and pediococci as versatile cell factories - Evaluation of strain properties and genetic tools. Biotechnol Adv. 35, 419–442.

Bringel, F., Castioni, A., Olukoya, D.K., Felis, G.E., Torriani, S. and Dellaglio, F. (2005) *Lactobacillus plantarum* subsp. *argentoratensis* subsp. nov., isolated from vegetable matrices. International Journal of Systematic and Evolutionary Microbiology. 55, 1629–34.

Brousseau, R., Hill, J.E., Prefontaine, G., Goh, S.H., Harel, J. and Hemmingsen, S.M. (2001) *Streptococcus suis* serotypes characterized by analysis of chaperonin 60 gene sequences. Applied and Environmental Microbiology. 67, 4828–33.

Carlier, J.P., Bonne, I. and Bedora-Faure, M. (2006) Isolation from canned foods of a novel *Thermoanaerobacter* species phylogenetically related to *Thermoanaerobacter mathranii* (Larsen 1997): Emendation of the species description and proposal of *Thermoanaerobacter mathranii* subsp. *Alimentarius* subsp. nov. Anaerobe. 12, 153–9.

Chaban, B., Links, M.G., Paramel Jayaprakash, T., Wagner, E.C., Bourque, D.K., Lohn, Z., Albert, A.Y.K., van Schalkwyk, J., Reid, G., Hemmingsen, S.M., Hill, J.E. and Money, D.M. (2014) Characterization of the vaginal microbiota of healthy Canadian women through the menstrual cycle. Microbiome. 2, 23.

Claesson, M.J., van Sinderen, D. and O’Toole, P.W. (2008) *Lactobacillus* phylogenomics-- towards a reclassification of the genus. International Journal of Systematic and Evolutionary Microbiology. 58, 2945–54.

Damé-Teixeira, N., Ev, L.D., Bitello-Firmino, L., Soares, V.K., Dalalba, R.S., Rup, A.G., Maltz, M. and Parolo, C.C.F. (2020) Characterization of Lactobacilli isolated from carious dentin after selective caries removal and cavity sealing. Arch Oral Biol. 121, 104988.

Dumonceaux, T.J., Hill, J.E., Hemmingsen, S.M. and Van Kessel, A.G. (2006a) Characterization of intestinal microbiota and response to dietary virginiamycin supplementation in the broiler chicken. Applied and Environmental Microbiology. 72, 2815–2823.

Dumonceaux, T.J., Hill, J.E., Pelletier, C., Paice, M.G., Van Kessel, A.G. and Hemmingsen, S.M. (2006b) Molecular characterization of microbial communities in Canadian pulp and paper activated sludge and quantification of a novel *Thiothrix eikelboomii*-like bulking filament. Canadian Journal of Microbiology. 52, 494–500.

Freitas, A.C., Bocking, A., Hill, J.E., Money, D.M., Money, D., Bocking, A., Hemmingsen, S., Hill, J., Reid, G., Dumonceaux, T., Gloor, G., Links, M., O’Doherty, K., Tang, P., van Schalkwyk, J., Yudin, M., and the VOGUE Research Group (2018) Increased richness and diversity of the vaginal microbiota and spontaneous preterm birth. Microbiome. 6, 117.

Guindon, S., Dufayard, J.-F., Lefort, V., Anisimova, M., Hordijk, W. and Gascuel, O. (2010) New Algorithms and Methods to Estimate Maximum-Likelihood Phylogenies: Assessing the Performance of PhyML 3.0. Syst Biol. 59, 307–321.

Haakensen, M., Dobson, C.M., Hill, J.E. and Ziola, B. (2009) Reclassification of *Pediococcus dextrinicus* (Coster and White 1964) back 1978 (Approved Lists 1980) as *Lactobacillus dextrinicus* comb. nov., and emended description of the genus *Lactobacillus*. International Journal of Systematic and Evolutionary Microbiology. 59, 615–21.

Haakensen, M., Pittet, V. and Ziola, B. (2011) Reclassification of *Paralactobacillus selangorensis* Leisner et al. 2000 as *Lactobacillus selangorensis* comb. nov. Int J Syst Evol Microbiol. 61, 2979–2983.

Hatem, A., Bozdağ, D., Toland, A.E. and Çatalyürek, Ü.V. (2013) Benchmarking short sequence mapping tools. BMC Bioinformatics. 14, 184.

Heo, J., Kim, S.-J., Kim, J.-S., Hong, S.-B. and Kwon, S.-W. (2020) Comparative genomics of Lactobacillus species as bee symbionts and description of Lactobacillus bombintestini sp. nov., isolated from the gut of Bombus ignitus. J Microbiol. 58, 445–455.

Hill, J.E. and Albert, A.Y.K. (2019) Resolution and co-occurrence patterns of *Gardnerella leopoldii, Gardnerella swidsinskii, Gardnerella piotii* and *Gardnerella vaginalis* within the vaginal microbiome. Infection and Immunity. 87, e00532–19.

Hill, J.E., Paccagnella, A., Law, K., Melito, P.L., Woodward, D.L., Price, D.L., Ng, L.-K., Hemmingsen, S.M. and Goh, S.H. (2006a) Identification of *Campylobacter* spp. and discrimination from *Helicobacter* and *Arcobacter* spp. by direct sequencing of PCR-amplified cpn60 sequences and comparison to cpnDB, a chaperonin reference sequence database. Journal of Medical Microbiology. 55, 393–399.

Hill, J.E., Town, J.R. and Hemmingsen, S.M. (2006b) Improved template representation in *cpn*60 PCR product libraries generated from complex templates by application of a specific mixture of PCR primers. Environmental Microbiology. 8, 741–746.

Hiu, S.F., Holt, R.A., Sriranganathan, N., Seidler, R.J. and Fryer, J.L. (1984) *Lactobacillus piscicola,* a new species from salmonid fish. International Journal of Systematic Bacteriology. 34, 393–400.

King, W.L., Siboni, N., Kahlke, T., Green, T.J., Labbate, M. and Seymour, J.R. (2019) A new high throughput sequencing assay for characterizing the diversity of natural *Vibrio* communities and Its application to a Pacific oyster mortality event. Front Microbiol. 10.

Koirala, R., Taverniti, V., Balzaretti, S., Ricci, G., Fortina, M.G. and Guglielmetti, S. (2015) Melting curve analysis of a groEL PCR fragment for the rapid genotyping of strains belonging to the *Lactobacillus casei* group of species. Microbiol Res. 173, 50–58.

Lefort, V., Longueville, J.-E. and Gascuel, O. (2017) SMS: Smart Model Selection in PhyML. Mol Biol Evol. 34, 2422–2424.

Li, H. (2018) Minimap2: pairwise alignment for nucleotide sequences. Bioinformatics. 34, 3094–3100.

Links, M.G., Dumonceaux, T.J., Hemmingsen, S.M. and Hill, J.E. (2012) The chaperonin-60 universal target is a barcode for bacteria that enables *de novo* assembly of metagenomic sequence data. PLoS ONE. 7, e49755.

Liu, D.D. and Gu, C.T. (2020) Proposal to reclassify *Lactobacillus zhaodongensis, Lactobacillus zeae, Lactobacillus argentoratensis* and *Lactobacillus buchneri* subsp. *silagei* as *Lacticaseibacillus zhaodongensis* comb. nov., *Lacticaseibacillus zeae* comb. nov., *Lactiplantibacillus argentoratensis* comb. nov. and *Lentilactobacillus buchneri* subsp. *silagei* comb. nov., respectively and *Apilactobacillus kosoi* as a later heterotypic synonym of *Apilactobacillus micheneri*. Int J Syst Evol Microbiol. 70, 6414–6417.

Minana-Galbis, D., Urbizu-Serrano, A., Farfan, M., Fuste, M.C. and Loren, J.G. (2009) Phylogenetic analysis and identification of *Aeromonas* species based on sequencing of the cpn60 universal target. International Journal of Systematic and Evolutionary Microbiology. 59, 1976–1983.

Muirhead, K., Perez-Lopez, E., Bahder, B.W., Hill, J. and Dumonceaux, T. (2019) The CpnClassiPhyR is a resource for cpn60 universal target-based classification of phytoplasmas. Plant Disease.

Oliver, K.L., Hamelin, R.C. and Hintz, W.E. (2008) Effects of transgenic hybrid aspen overexpressing polyphenol oxidase on rhizosphere diversity. Appl Environ Microbiol. 74, 5340–8.

Pantucek, R., Sedlacek, I., Petras, P., Koukalova, D., Svec, P., Stetina, V., Vancanneyt, M., Chrastinova, L., Vokurkova, J., Ruzickova, V., Doskar, J., Swings, J. and Hajek, V. (2005) *Staphylococcus simiae* sp. nov., isolated from South American squirrel monkeys. International Journal of Systematic and Evolutionary Microbiology. 55, 1953–8.

Parte, A.C., Sardà Carbasse, J., Meier-Kolthoff, J.P., Reimer, L.C. and Göker, M. (2020) List of Prokaryotic names with Standing in Nomenclature (LPSN) moves to the DSMZ. International Journal of Systematic and Evolutionary Microbiology,. 70, 5607–5612.

Pearson, W.R. and Lipman, D.J. (1988) Improved tools for biological sequence comparison. Proceedings of the National Academy of Sciences of the United States of America. 85, 2444–8.

Pratt, D.L., Dumonceaux, T.J., Links, M.G. and Fonstad, T.A. (2012) Influence of mass burial of animal carcasses on the types and quantities of microorganisms within a burial site. Transactions of the American Society of Agricultural and Biological Engineers. 55, 2195–2212.

Rice, P., Longden, I. and Bleasby, A. (2000) EMBOSS: the European Molecular Biology Open Software Suite. Trends in Genetics. 16, 276–7.

Sakamoto, M., Ikeyama, N., Kunihiro, T., Iino, T., Yuki, M. and Ohkuma, M. (2018) *Mesosutterella multiformis* gen. nov., sp. nov., a member of the family *Sutterellaceae* and *Sutterella megalosphaeroides* sp. nov., isolated from human faeces. International Journal of Systematic and Evolutionary Microbiology. 68, 3942–3950.

Sakamoto, M. and Ohkuma, M. (2010) Usefulness of the hsp60 gene for the identification and classification of Gram-negative anaerobic rods. Journal of Medical Microbiology. 59, 1293–302.

Sakamoto, M., Suzuki, N. and Benno, Y. (2010) hsp60 and 16S rRNA gene sequence relationships among species of the genus *Bacteroides* with the finding that *Bacteroides suis* and *Bacteroides tectus* are heterotypic synonyms of *Bacteroides pyogenes*. International Journal of Systematic and Evolutionary Microbiology. 60, 2984–90.

Sauer, M., Russmayer, H., Grabherr, R., Peterbauer, C.K. and Marx, H. (2017) The Efficient Clade: Lactic Acid Bacteria for Industrial Chemical Production. Trends Biotechnol. 35, 756–769.

Schellenberg, J.J., Paramel Jayaprakash, T., Withana Gamage, N., Patterson, M.H., Vaneechoutte, M. and Hill, J.E. (2016) *Gardnerella vaginalis* subgroups defined by cpn60 sequencing and sialidase activity in isolates from Canada, Belgium and Kenya. PLoS One. 11, e0146510.

Vancuren, S.J. and Hill, J.E. (2019) Update on cpnDB: a reference database of chaperonin sequences. Database (Oxford). 2019.

Vancuren, S.J., Santos, S.J.D., Hill, J.E. and Team, the M.M.L.P. (2020) Evaluation of variant calling for cpn60 barcode sequence-based microbiome profiling. PLOS ONE. 15, e0235682.

Verbeke, T.J., Sparling, R., Hill, J.E., Links, M.G., Levin, D. and Dumonceaux, T.J. (2011) Predicting relatedness of bacterial genomes using the chaperonin-60 universal target (*cpn*60 UT): application to *Thermoanaerobacter* species. Systematic and Applied Microbiology. 34, 171–179.

Xie, M., Pan, M., Jiang, Y., Liu, X., Lu, W., Zhao, J., Zhang, H. and Chen, W. (2019) groEL gene-based phylogenetic analysis of *Lactobacillus* species by high-throughput sequencing. Genes (Basel). 10.

Yang, B., Chen, Y., Stanton, C., Ross, R.P., Lee, Y.-K., Zhao, J., Zhang, H. and Chen, W. (2019) *Bifidobacterium* and *Lactobacillus* composition at species level and gut microbiota diversity in infants before 6 weeks. Int J Mol Sci. 20.

Zheng, J., Wittouck, S., Salvetti, E., Franz, C.M.A.P., Harris, H.M.B., Mattarelli, P., O’Toole, P.W., Pot, B., Vandamme, P., Walter, J., Watanabe, K., Wuyts, S., Felis, G.E., Gänzle, M.G. and Lebeer, S. (2020) A taxonomic note on the genus *Lactobacillus*: Description of 23 novel genera, emended description of the genus *Lactobacillus* Beijerinck 1901, and union of Lactobacillaceae and Leuconostocaceae. International Journal of Systematic and Evolutionary Microbiology. 70, 2782–2858.

